# Microridges are apical projections formed of branched F-actin networks that organize the glycan layer

**DOI:** 10.1101/442871

**Authors:** Clyde Savio Pinto, Ameya Khandekar, Rajasekaran Bhavna, Petra Kiesel, Gaia Pigino, Mahendra Sonawane

## Abstract

Apical projections are integral functional units of epithelial cells. Microvilli and stereocilia are cylindrical apical projections that are formed of bundled actin. Microridges on the other hand, grow laterally long, forming labyrinthine patterns on surfaces of various kinds of squamous epithelial cells. So far, the structural organization and functions of microridges have remained elusive. We have analyzed microridges on zebrafish epidermal cells using confocal and electron microscopy methods including electron tomography, to show that a microridge is formed of a network of F-actin and requires the function of the Arp2/3 complex for its maintenance. During development, microridges begin as F-actin punctae showing signatures of branching and requiring an active Arp2/3 complex. Using inhibitors of actin polymerization and the Arp2/3 complex, we show that microridges organize the surface glycan layer. Our analyses have unraveled the F-actin organization supporting the most abundant and evolutionarily conserved apical projection, which functions in glycan organization.

## Introduction

Epithelial tissues cover the outer surface of metazoans as well as the outer and inner surfaces of organs, thus separating two physiologically distinct compartments. Over the course of evolution, epithelia have undergone diversification and specialization allowing them to adapt to various conditions and perform not only the barrier function but also additional functions such as absorption, secretion and mechanosensation (*Le Bivic*, 2013; *Urdy et al.*, 2016). These functions require special adaptions of their apical surfaces (*Apodaca and Gallo*, 2013; *Rodriguez-Boulan and Nelson*, 1989), which include diverse apical membrane protrusions. Additionally, the apical surface is kept hydrated and protected from pathogens and physical injury by the overlying glycocalyx and mucosal layers (*Apodaca and Gallo*, 2013; *Cone*, 2009). Aside from the glycocalyx, mucosal epithelia have mucus - a viscoelastic fluid - on their surface. Thus, the apical zone of epithelial tissues is comprised of three components, namely, the apical plasma membrane with its projections, the cytoskeleton supporting these projections and the outer mucus-glycocalyx layer (*Apodaca and Gallo*, 2013).

In vertebrates, epithelial cells exhibit membrane protrusions of various shapes and sizes. Columnar epithelia possess actin-based cylindrical microvilli or their derivatives such as stereocilia (*Apodaca and Gallo*, 2013; *Sauvanet et al.*, 2015; *Tilney et al.*, 1980). On the contrary, the apical domain of several of the non-cornified squamous epithelia is thrown into laterally long protrusions called microplicae or microridges (*Apodaca and Gallo*, 2013). While previous studies have significantly enhanced our understanding of actin organization in microvilli (*Crawley et al.*, 2014; *Sauvanet et al.*, 2015), how actin is organized to build microridges remains controversial.

Mature microridges are relatively stable wall like protrusions, arranged in a labyrinthine fashion (*Bereiter-Hahn et al.*, 1979; *Collin and Collin*, 2000; *Fishelson*, 1984; *Hawkes*, 1974; *Lam et al.*, 2015; *Schliwa*, 1975). They are shown to be present on several non-cornified epithelia in various species across vertebrate phyla, making them one of the more widespread actin protrusions (*Bereiter-Hahn et al.*, 1979; *Breipohl et al.*, 1977; *Collin and Collin*, 2000; *Eroschenko and Osman*, 1986; *Fenger and Knoth*, 1981; *Hafez et al.*, 1977; *Johnston et al.*, 1996; *Makabe et al.*, 1998; *Saito and Itoh*, 1993; *Uehara et al.*, 1991; *Worawongvasu*, 2007). Microridges are thought to have important roles in mucus retention, abrasion resistance and increasing the tensile strength of the apical domain (*Fishelson*, 1984; *Sharma et al.*, 2005; *Sperry and Wassersug*, 1976). They have also been proposed to function as a rapidly deployable store of F-actin and membrane during wound healing (*Sharma et al.*, 2005).

The nature of the protrusion is determined by the organization and dynamics of F-actin within it. Filaments in actin protrusions take on one of two conformations - either as a branched network of filaments, seen in lamellipodia (*Svitkina and Borisy*, 1999) or as parallel bundles forming the core of projections like microvilli and stereocilia (*Revenu et al.*, 2004). Although the importance of the Arp2/3 complex, an important regulator of branched actin networks, in the maintenance of microridges in zebrafish epidermal cells is known (*Lam et al.*, 2015), it is not clear how actin within them is organized. While actin within microridges of guppy epidermal cells is reported as bundled (*Bereiter-Hahn et al.*, 1979), microridges on carp oral mucosal cells, cat epithelial cells, and guppy cells in culture are shown to contain a network of actin (*Andrews*, 1976; *Bereiter-Hahn et al.*, 1979; *Uehara et al.*, 1991). Furthermore, based on scanning ion-conductance microscopy, it has been proposed that microvilli can merge to form ridges (*Gorelik et al.*, 2003), suggesting that microridges form from microvilli containing F-actin bundled cores. The zebrafish periderm offers a genetically and microscopically tractable model to study the formation, structure and function of microridges. It has been recently shown that the cell polarity regulator atypical Protein Kinase C (aPKC) plays a role in restricting the elongation of microridges by controlling levels of Lethal giant larvae (Lgl) and nonmuscle MyosinII at the apical domain of peridermal cells (*Raman et al.*, 2016). However, the ultrastructure and function of these protrusions have not been characterized well. Here, we have performed molecular as well as structural analysis of microridges in the developing zebrafish epidermis and probed for their functional importance in the context of the mucus and glycocalyx, which we collectively call the glycan layer. We show that the molecular composition of microridges is similar to lamellipodia and that Arp2/3 function is essential for their formation and maintenance. Electron tomography (ET) followed by segmentation analysis reveals that the actin is organized in the form of a network. We further disprove the notion that microridges are formed from microvilli by showing that small actin spots that generate microridges consist of the Arp2/3 complex. Lastly, our analyses also indicate that microridges are important for the organization of the glycan layer.

## Results

### Ultrastructural characterization of the apical zone of peridermal cells

The ultrastructural characterization of the entire apical zone including the microridge and its surroundings has not been carried out so far in the developing zebrafish periderm. We looked at the organization of the peridermal cell in transmission electron micrographs (TEM) as well as in scanning electron micrographs (SEM) at 48 hpf — a time point at which embryos begin to hatch and presumably require functional microridges.

TEM analysis using routine glutaraldehyde (GA) and osmium tetroxide fixation revealed three distinct zones in the peridermal cell- the protrusive zone consisting of microridges, the sub-protrusive zone comprised of actin and keratin filaments and a basal region containing various organelles (Figure 1 A). Microridges had a mean height of 315±82 nm (n= 245, N=3; Fig. S 1 B), as measured from TEM. Using SEM (Fig. S 1 A), we found their mean width to be 162±36 nm (n= 645, N=3; Fig. S 1 B). To observe the organization of the glycan layer, we fixed samples in the presence of alcian blue and lysine, which has been shown to preserve the glycan layer better (*Fassel et al.*, 1997, 1992; *Fischer et al.*, 2012). SEM of heads of animals fixed in such a way showed that the peridermal surface was covered by a thick glycan layer, which completely obscures the underlying microridges on the head epidermis (Figure 1 B). TEM analysis of such a sample revealed glycan enrichment around the microridges (Fig. S 1 C).

**Figure 1.**
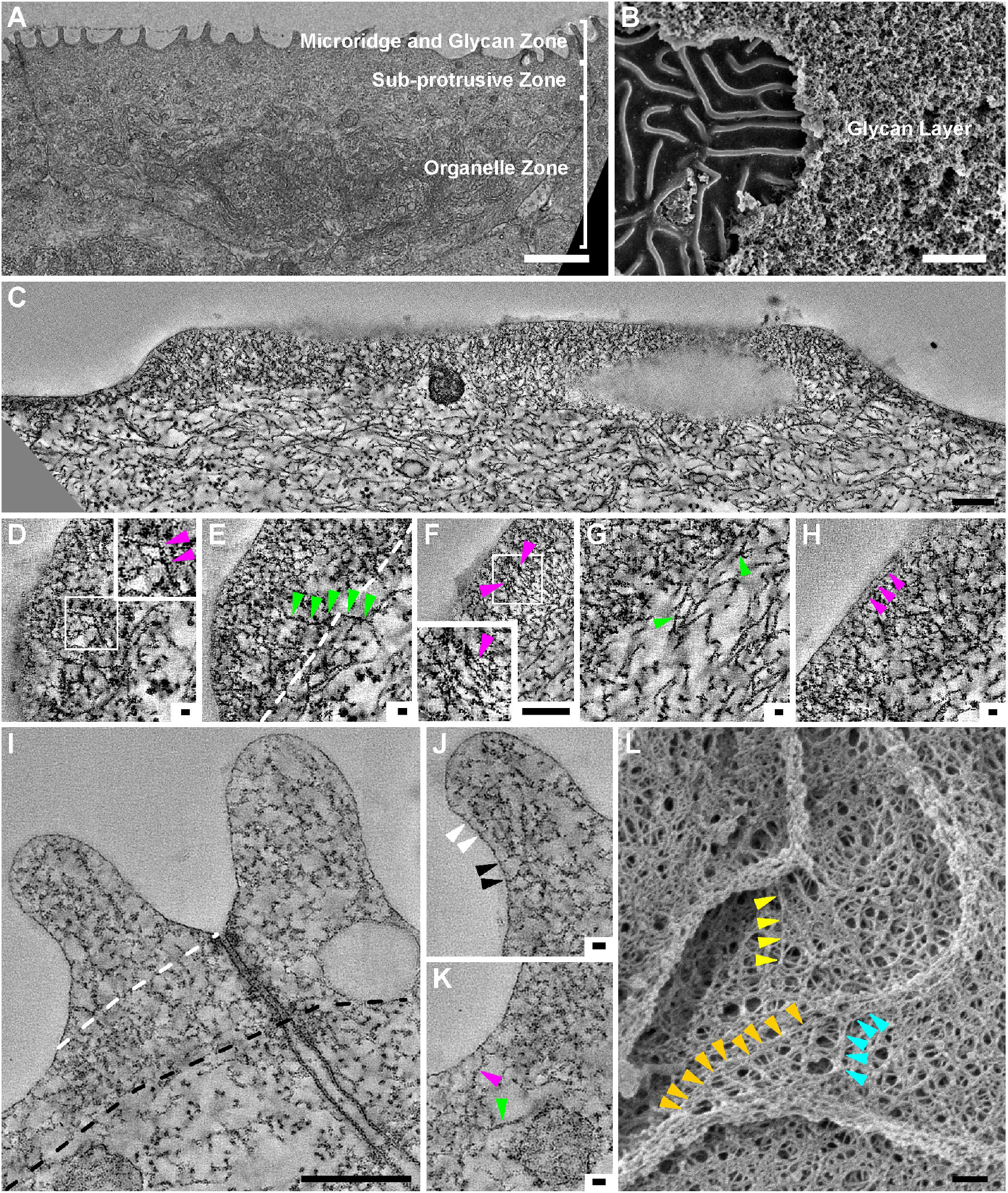
EM tomography of untreated samples and SEM on detergent extracted samples reveals the ultrastructural organization of the actin microridge, sub-protrusive zone and cortex. TEM through a head peridermal cell revealed that peridermal cells comprise of a microridge/glycan zone, sub-protrusive zone and organelle zone (A). SEM of an Alcian blue and lysine fixed head periderm sample (B) reveals a thick glycan layer, which is chipped off on the left revealing the underlying microridges. Tomograms of Gluteraldehyde+OsO_4_ fixed samples through the length of the microridge (C) and perpendicular to the cell-boundary microridge (I). Zoomed in areas of C and I are shown in (D-H) and (J,K), respectively. Each image in C-H and I-K represents a 0.7 and 0.55 nm thick slice of the tomogram in the Z axis respectively. (L) SEM of a detergent extracted sample. Magenta and green arrowheads indicate actin and keratin filaments, respectively. In the inset in (D), the arrowheads point to a filament with branch points. In (E) the arrowheads point to a keratin filament entering the microridge from the sub-protrusive zone whereas the white dashed line demarcates the boundary between the microridge above from the sub-protrusive zone below. Small filaments that appear to be parallel to each other (F) are shown by the arrowheads and inset in F. The arrowheads in (G) point to branch points in keratin filaments in the sub-protrusive zone. In (H) the arrowheads point to a filament that is parallel to the plasma membrane. The cytoskeleton from the top of the microridge to its base (indicated by white dashed line in I) and the sub-protrusive zone till the base of the tight junction (indicated by the black dashed line) is denser and made up largely of actin. In (J) smaller actin membrane crosslinks (white arrowhead) and longer actin membrane crosslinks (black arrowhead) are shown. The cortex between ridges (K) is made up largely of actin, which are thinner filaments (magenta arrowhead), distinctly different from the thicker keratin filament below (green arrowhead). Coloured arrowheads in (L) indicate filaments or filament bundles extending from the cortex into the microridge. The scale bars in A,B are 2 μm, in C,F,I,L are equal to 200 nm and those in D,E,G,H,J,K are 20 nm.

Since conventional TEM did not reveal the F-actin organization within the microridge due to the dense packing of actin (Fig. S 1 D), we used Electron Tomography (ET) to unravel the structural details of actin organization (Figure 1 C-K). We produced tomograms (Videos 1-3), with different orientations of microridges. Samples fixed using osmium and GA (Figure 1 C-K; Video 1; Video 2) showed that filaments within a microridge were largely arranged at different angles to each other (Figure 1 C-J), suggestive of a network of actin along with a few parallel filaments (Figure 1 F). We used an algorithm (see methods) to segment filaments in a tomogram of microridges (Figure 2). This allowed a reconstruction of the filaments within the microridge as shown, within a small region of the tomogram (Figure 2 B-F, Video 4). These corroborated the fact that filaments are organized in a network like fashion (Figure 2 E,F; Video 4). We were also able to observe branching of filaments (Figure 1 D; Figure 2 G,H). ET of samples fixed using tannic acid and uranyl acetate, avoiding OsO_4_, corroborated the fact that actin within the microridge is organized as a network (Video 3). Aside from a few filaments parallel to the microridge membrane (Figure 1 H), we observed a number of actin-membrane crosslinks at the interphase between the membrane and the actin network (Figure 1 J). Interestingly, there were many instances in which vesicles were found in the microridge, particularly at its base, indicating that it is not an isolated system like a cilium (Figure 1 I; Fig. S 1 E).

**Figure 2.**
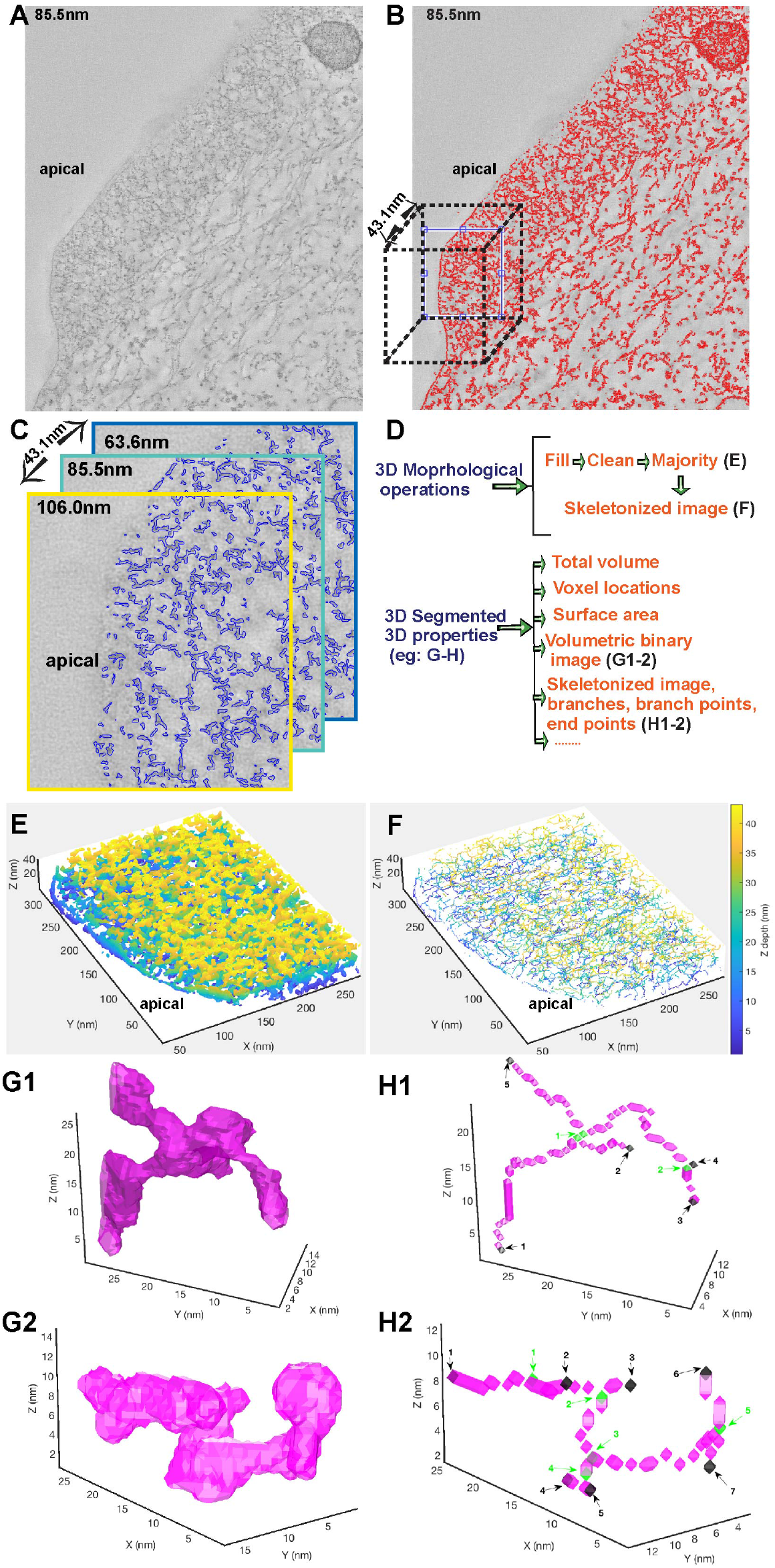
Segmentation of an electron tomogram reveals the arrangement of actin to be a network within the microridge. An EM image section tomogram was taken with 2090×1740×249 voxels at an equal spacing of 0.707nm in each direction, here shown at a depth of 85.5nm (A). Higher Eigen value 2D matrix outlines of the actin structures within the microridges shown in (B). A cubic section of 283.5×318.8×43.1 nm3 (dotted cube in B) is cropped for illustration of the segmentation analysis. Image binarization outlines (in blue) of the actin structures (C), shown at 3 different depths. (D) Steps showing subsequent 3D morphological operations on the reconstructed 3D binary stack followed by computation of segmented structures properties. 3D reconstruction of actin (E) and corresponding 3D skeleton image (F) reveal the arrangement of actin meshwork. Depth is indicated by a colorbar. Examples of single rendered actin structures in magenta (G1-2) and corresponding skeletons (H1-2) within the microridges indicate branching of filaments. Branch points (green) and endpoints (black) are highlighted on the 3D skeleton images. All the branchpoints and endpoints are numbered and indicated by arrow marks. The EM tomogram analysed here was a part of the tomogram shown in Figure 1 C-H and Video 1.

ET revealed that the sub-protrusive zone contained cortical actin and a network of thick keratin filaments with discernible branching points (Figure 1 C,G,I,K). Some keratin filaments —discernable by their thickness as compared to actin (Figure 1 K) — clearly entered the main body of the microridge, indicating that keratin and actin filaments arising from the cortex support its architecture (Figure 1E). To confirm such a contribution from the cortex, we used detergent extraction to remove apical membranes of peridermal cells and performed SEM. In these preparations, actin filaments in the cortex were clearly visible (Figure 1 L). These filaments were arranged in a web like pattern. Interestingly, we found that actin filaments and filament bundles extended a considerable distance within the cortex and into the microridge (Figure 1 L). Further corroborating with these ET and SEM data, we observed the localization of keratin - using the AE1/AE3 antibody - to the apical cortex and occasionally in the microridge (Fig. S 2A). We did not observe any direct association of microtubules with the microridge either by ET or immunostaining using an *α*-tubulin antibody (Fig. S 2 B).

To conclude, the apical zone of zebrafish peridermal cells consists of a glycan layer, an apical domain consisting of short projections called microridges and the cytoskeleton that supports them. Within the microridge, F-actin is organized in the form of a network, supported by filaments arising from the sub-protrusive zone or terminal web and cortex. The interior of the microridge space is accessible to vesicles indicating that the microridge is not an isolated system.

### Molecular signature of the microridge in zebrafish

The microridge is strikingly different in both morphology and F-actin organization as compared to microvilli (*Crawley et al*., 2014; *Sauvanet et al*., 2015). This prompted us to look at the molecular composition of microridges. Thus far, only a few actin-binding proteins are known to localize to the microridge. These include *α*-actinin, ezrin, myosin II, VASP and cortactin (*Bereiter-Hahn et al.*, 1979; *DePasquale*, 2018; *Lam et al.*, 2015; *Raman et al.*, 2016). Since actin-binding proteins regulate and organize actin filaments into specific arrangements such as parallel or antiparallel bundles and branched networks, we analyzed the localization of additional regulators of the actin cytoskeleton to microridges. We utilized two candidate-based strategies — a) antibody stainings to identify proteins endogenously present at the microridge and b) localization of plasmid encoded expression of tagged proteins. Our immunolocalization analysis revealed that ArpC2, a component of the actin nucleator Arp2/3 complex, localizes to microridges at 48 hpf (Figure 3 A). The nucleation-promoting-factor WASp-like (WASL) (Figure 3 B) and Cofilin, an actin severing and depolymerizing protein, were also found to localize to the microridge (Figure 3 C). Our analysis using various tagged proteins revealed that the actin bundling and crosslinking proteins Eplin/Lima-1, found in stress fibers or at the adherens junction (*Abe and Takeichi*, 2008; *Maul et al.*, 2003), and Filamin, found in branched structures like lamellipodia or membrane ruffles (*Small et al.*, 2002; *Vadlamudi et al.*, 2002), also localized to microridges (Figure 3 F-K).

**Figure 3.**
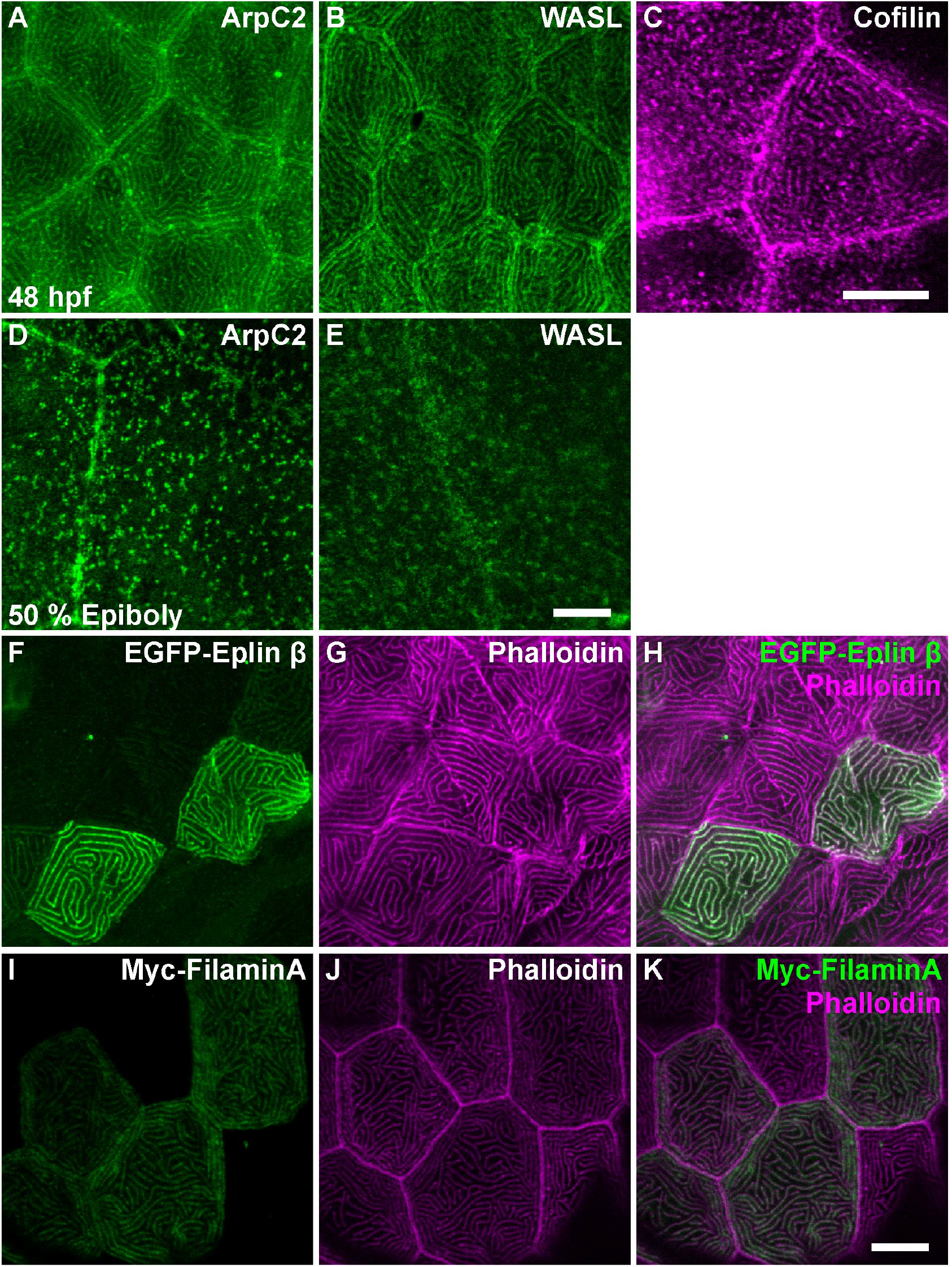
Localization of actin binding proteins at the microridge.Confocal micrographs showing immuno-localization of ArpC2 (A, D), WASL (B, E), Cofilin (C) at 48 hpf (A-C) and at 50% epiboly (D,E). Plasmid encoded expression of Eplin *β* (F), and Filamin A (I) in green localize to microridges marked with phalloidin (G,J) in magenta and respective merges (H,K). Plasmids were injected at the 1 cell stage and expressed in a clonal fashion. Scale bars in E for A,B,D,E, in C for C, in K for F-K are equal to 10 μm.

Since the activity of the Arp2/3 complex is essential for maintenance of microridges (*Lam et al.*, 2015), we asked whether there is a temporal correlation between the localization of the Arp2/3 complex, WASL and microridge formation. As the Arp2/3 complex and WASL are bona-fide components of branched actin, we also reasoned that such an analysis would allow us to test the notion whether microridges are formed from microvilli. As a prerequisite for this analysis, we characterized the development of microridges starting from 50% epiboly (Fig. S 3). We found that at 50% epiboly, F-actin punctae were present at the apical domain of EVL (Enveloping layer) cells (Fig. S 3 A), which later give rise to the periderm (*Kimmel et al.*, 1990). Such punctae were visible till 12 hpf (Fig. S 3 B,C), after which slightly long F-actin structures were observed near the cell periphery. These seem to further elongate over time (Fig. S 3 D-I). To check if these punctae were associated with protrusions, we preformed SEM. We observed small protrusions at the periphery but not the center of the cell in most parts of the EVL at 9 hpf (Fig. S 3 J). By 18 hpf, microridges at the cell boundary became prominent and by 48 hpf most of the peridermal cell surface was covered with microridges (Fig. S 3 K,L). Both the Arp2/3 complex and WASL showed punctate localization at the apical cortex of EVL cells as early as 50% epiboly, similar to F-actin punctae (Figure 3 D,E). Inhibition of the Arp2/3 complex by CK666 at 9 hpf resulted in a stark reduction in F-actin punctae (Figure 4 A-C), inhibition at 18 hpf prevented the formation of microridges (Figure 4 D-F) and at 47 hpf resulted in their breakdown (Figure 4 G-I), indicating that the Arp2/3 complex is important for both the formation and maintenance of microridges and punctae. Since Keratins contribute to the terminal web, we also investigated if a temporal link exists between the formation of the keratin cytoskeleton and microridges. Keratin levels, analyzed by AE1/AE3 antibody staining, were low at 9 hpf, increasing over time as monitored till 15 hpf (Fig. S 4).

**Figure 4.**
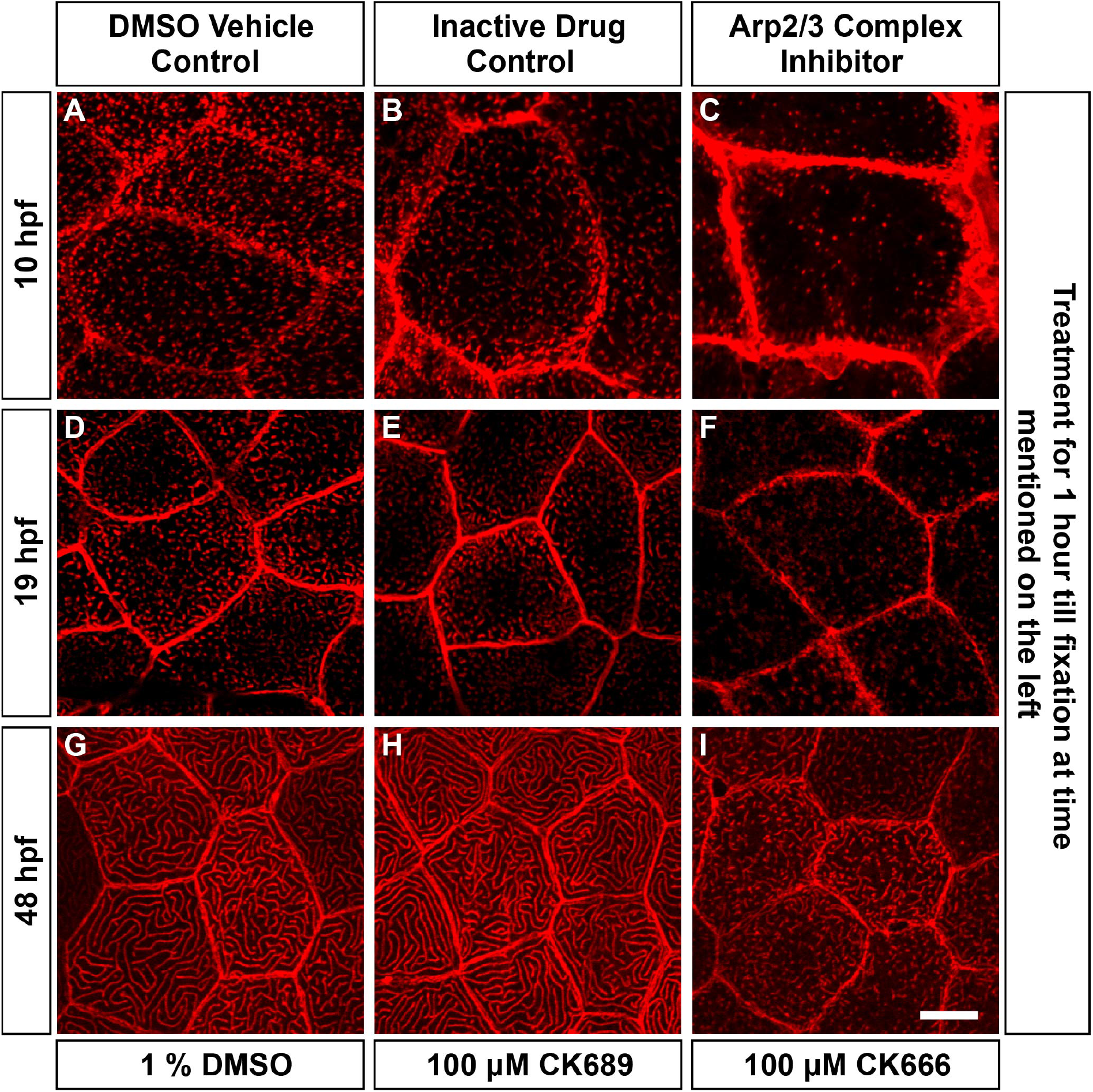
Arp2/3 complex activity is required for microridge formation and maintenance. Confocal microscopy analyses of phalloidin staining of embryos and larvae (A-I) treated with 1% DMSO (A,D,G), 100 μM CK689 (inactive drug control;B,E,H) and 100 μM CK666 (Arp2/3 complex inhibitor; C,F,I) for 1 hour from 9-10 hpf (A-C), 18-19 hpf (D-F) and 47-48 hpf (G-I). In all cases inhibition resulted in a breakdown of microridges or a decrease in phalloidin punctae relative to controls (N = 2 sets, n = 10). Scale bar in I for A-I is equivalent to 10 μm.

Thus, bona fide markers of branched actin networks such as the Arp2/3 complex, WASL and Filamin, as well as the actin-bundling protein Eplin, localize to microridges. The Arp2/3 complex is essential for the formation or maintenance of the actin punctae and microridges. At all time-points analyzed, actin structures contained the Arp2/3 complex, consequently microvilli-like structures were not observed during the time resolved experiments. Therefore, microridges form from actin punctae, presumably having branched configuration, and not from bundle based microvilli.

### Physiological relevance of microridges

The widespread occurrence of microridges on mucosal epithelial tissues has led researchers to propose that microridges function in retention and or distribution of mucus (*Fishelson*, 1984; *Schliwa*, 1975; *Sperry and Wassersug*, 1976). The developing zebrafish embryo is covered by a glycan layer as early as the epiboly stages (*Baskin et al.*, 2010), making it an excellent model to study the role of microridges in maintaining the organization of the glycan layer.

In order to visualize the glycan layer with a confocal microscope, we used a fluorescently tagged wheat germ agglutinin (WGA) lectin. This approach revealed that the glycan layer was arranged around microridges (Figure 5 A-C,E-G,I-K,M-O). To assess if microridges are important for the organization of the glycan layer, we used chemical inhibitors to disrupt the actin cytoskeleton and consequently the microridge pattern. While in control embryos, the WGA fluorescence followed the microridge pattern (Figure 5 A-C), in the CK666 treated embryos the WGA fluorescence was comparatively more uniform suggesting a mild effect on the organization of the glycan layer (Figure 5 E-G). SEM analysis on CK666 treated samples did not reveal a major change in glycan organization (Figure 5 D,H), again pointing to the mild effect of CK666 on the glycan layer (Figure 5 D,H). We achieved a severe perturbation of microridges using the actin monomer binding drug Latrunculin A (Lat A). LatA treatment resulted in the decreased density and length of microridges (Figure 5 I-P). We analysed the ventral region of the flank where the effect of LatA on microridges was most severe. As compared to the control (Figure 5 I-K), the LatA treated samples showed more intense WGA staining around the microridge remnants indicating accumulation of glycans around them (Figure 5 M-O). SEM analysis revealed that the glycan layer persisted in the troughs between microridges at a lower density. However, it was clearly enriched around the microridge remnants (Figure 5 L,P).

**Figure 5.**
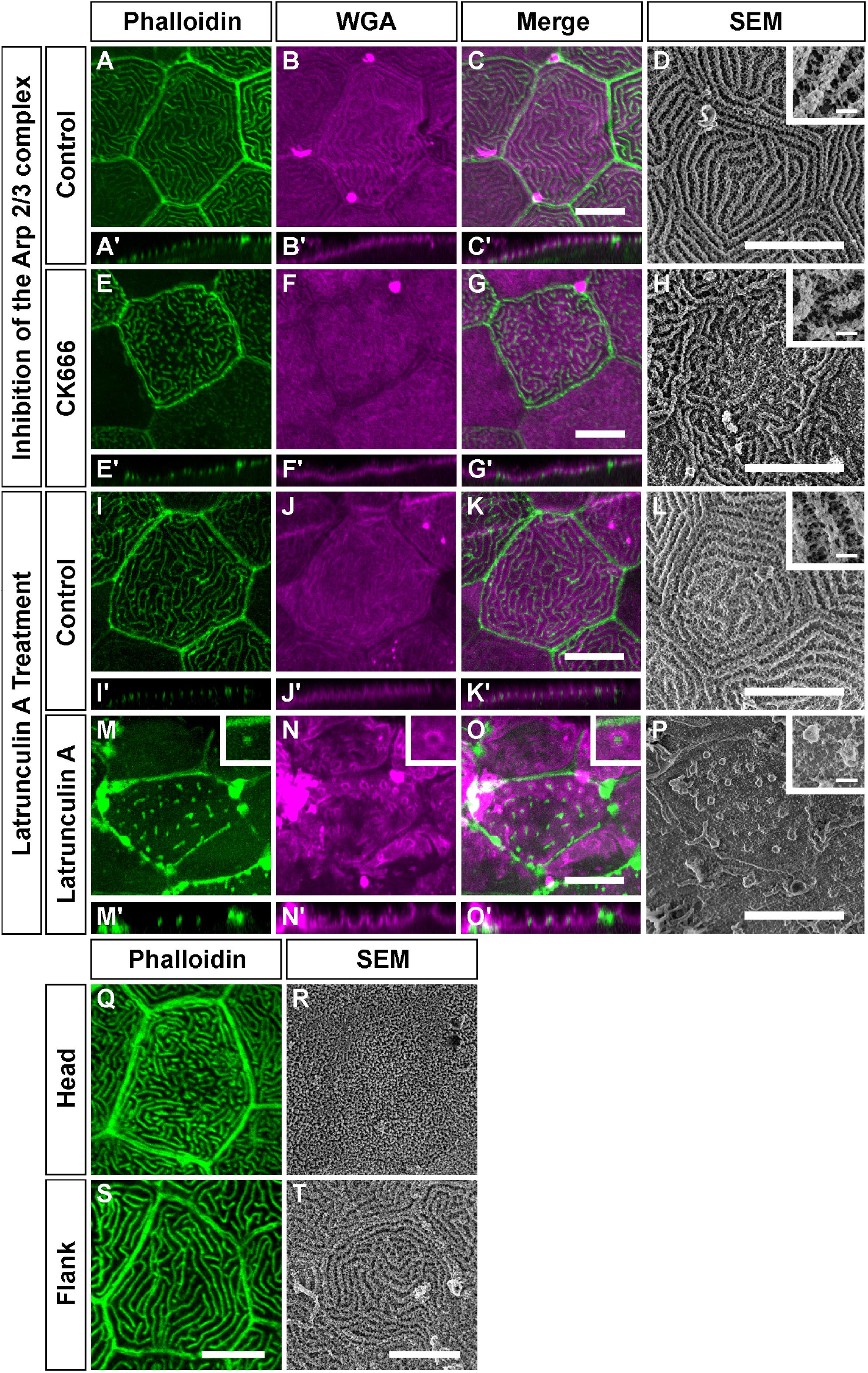
Microridges play a role in the organization of the glycan layer. Effect of the inhibition of the Arp2/3 complex on the glycan layer (A-H). Confocal micrographs of Phalloidin (A,E), WGA stainings (B,F), Merges (C, G) and SEM (D,H) of animals treated with 1% DMSO as vehicle control (A-D) or 100 μM CK666 (E-H) and respective orthogonal sections below the images (A’- C’,E’-G’). Effect of Latrunculin A treatment on the glycan layer (I-P). Phalloidin (I,M), WGA stainings (J,N) and Merges (K,O) of animals treated with 1% DMSO as vehicle control (I-K) or 2 μM Latrunculin A (M-O) and respective orthogonal sections below the images (M’- O’). Insets in M-O show a microridge remnant (phalloidin, green) surrounded by the glycan layer (WGA staining, magenta). SEM using Alcian blue and Lysine fixation of 1% DMSO as vehicle control (L) or 2 μM Latrunculin A (P). The insets in D,H,L,P show higher magnification SEM images of their respective treatments. Differences in the glycan layer organization in different parts of the animal (Q-T). Phalloidin staining (Q, S) and SEM (R, T) of the head (Q,R) and flank (S,T) peridermal cells. Note that microridges on the head are relatively densely packed with a flat glycan layer as compared to the flank. Scale bars in C for A-C, in G for E-G, in K for I-K, in O for M-O, in S for Q,S, in T for R,T and in D,H,L,P are 10 μm and in the insets in D,H,L,P are 1 μm.

During our analyses, we observed that microridge patterns differed on the flank (Figure 5 S,T) from those on the head (Figure 5 Q,R), with a higher packing density of microridges on the head. SEM analysis revealed that the glycan layer coats microridges on the head in a sheet like fashion without revealing their underlying pattern (Figure 5 R). In contrast, the glycan layer on the flank follows the pattern of underlying microridges, with glycans enriched on microridges with apparent gaps in between (Figure 5 T). This further indicates that microridges have the ability to retain glycans in their close proximity.

In conclusion, peridermal cells on the zebrafish larval epidermis are covered with a thick glycan layer. The organization of this layer relies on the underlying microridges.

## Discussion

The apical surface is functionally the most important part of an epithelial tissue. It is decorated with actin based projections and a glycan layer. Laterally long protrusions such as microridges have been known to morphologists for over 50 years now but have not been well studied. The zebrafish periderm, which is completely covered with microridges, offers an ideal system to study such protrusions and their role in glycan biology (*Lam et al.*, 2015; *Raman et al.*, 2016). Here, we used immunofluorescence, EM tomography and SEM analysis to understand how actin microridges form, are maintained and whether they function in glycan organization.

Our ultra-structural analyses presented here indicate that the microridge is largely made up of a network of actin. This organization is in fact similar to the one in lamellipodia, wherein the lamellipodial network meets the transverse actin arc (*Burnette et al.*, 2011). Our results, in zebrafish larvae at 48 hpf are consistent with Uehara et al, who have shown that microridges in carp oral mucosa exhibit actin filaments oriented randomly (*Uehara et al.*, 1991). However, unlike our ET analysis, this earlier analysis was done with detergent extracted samples and hence did not reflect the complexity of the network in its entirety. Our structural analysis is largely in agreement with the fact that all the major regulators of branched actin conformation - the Arp2/3 complex, WASL, Cofilin and Filamin - localize to the microridges and that Arp2/3 complex function is required in forming and maintaining microridges. In this sense, microridges can be thought of as several stable lamellipodium like protrusions organized in a labyrinthine manner on the apical surface of cells (Figure 6 A-C). Both SEM and tomographic data show that the microridge cytoskeleton is contiguous with the cortex and keratin network, suggesting that the organization of this layer would influence the microridge pattern (Figure 6 B). Curiously, in lamellipodia the sub-protrusive zone is formed of only actin, whereas in microridges it is formed of both actin and keratin, similar to microvilli (Figure 6 B,D,E). In fact, keratin filaments from the sub-protrusive zone also enter microridges, possibly offering additional structural support (Figure 6 B).

**Figure 6.**
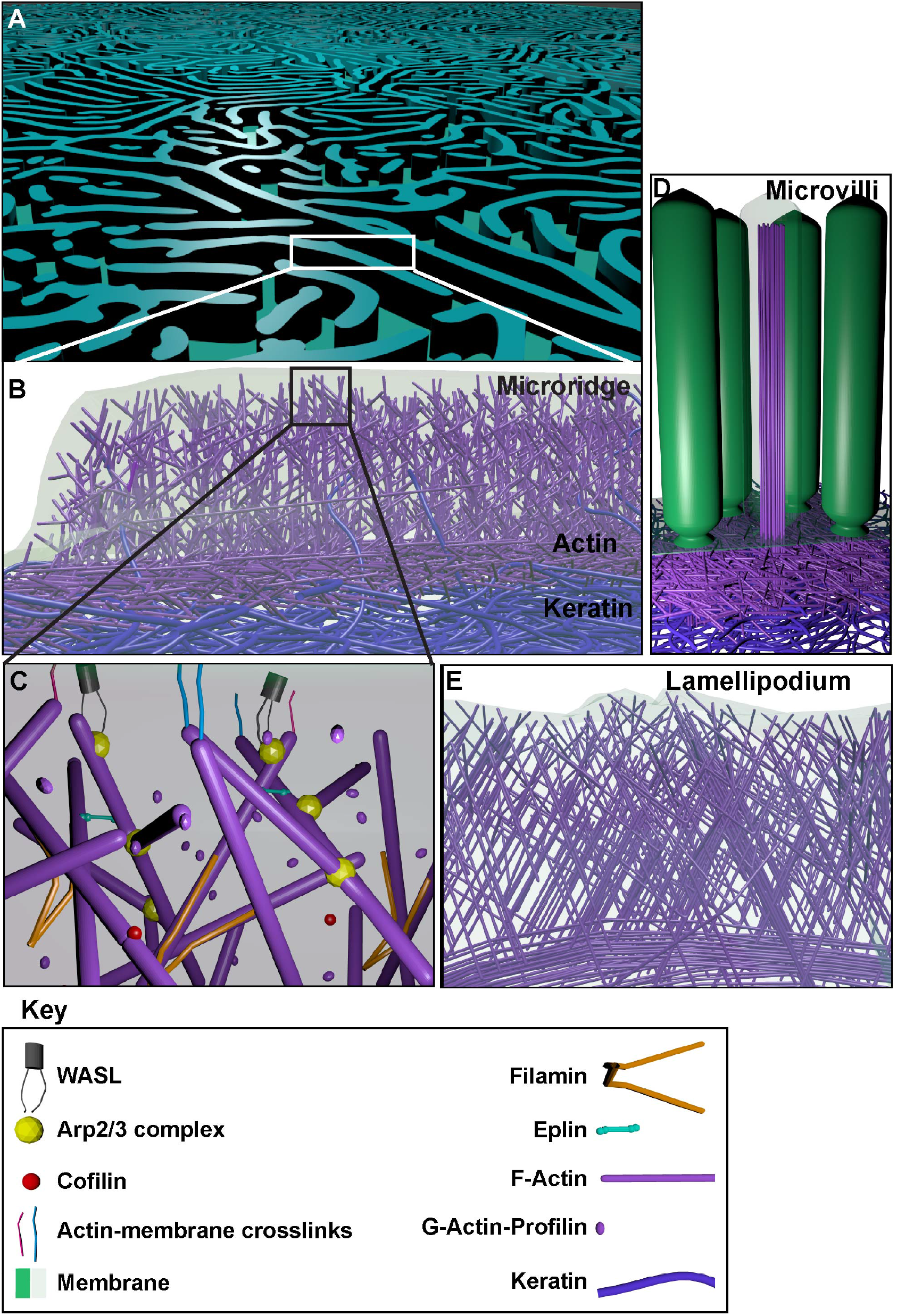
A schematic representation of the actin microridge based on EM and immunolocalization analyses. The actin microridge is a laterally long protrusion (A). It consists of a branched network of actin with an underlying sub-protrusive zone of actin (magenta) and keratin (blue). Occasionally, keratin filaments also enter the protrusion (B). The actin microridge protrusion itself consists of various molecules (C), many of which associate with branched networks The sub-protrusive zone of the actin microridge resembles the sub-protrusive terminal web of brush border microvilli (D), which are protrusions made up of bundled actin with a terminal web of actin and keratin. On the other hand, the branched actin architecture of the microridge protrusion has similarities with the lamellipodial protrusion (E), though lamellipodia have an actin based sub-protrusive zone, without a noticeable keratin cytoskeleton.

Intriguingly, we also observed localization of Eplin, a component of adherens junctions, as well as Fascin (Khandekar and Sonawane, unpublished observations), a component of bundled actin, to microridges. It is possible that though the F-actin network supporting microridges is largely controlled by regulators of branched actin, there might be a small but significant contribution from regulators of bundled actin. This is not surprising given the fact that structures formed of bundled actin, such as microspikes and filopodia, are associated with lamellipodia. Alternatively, such components of bundled actin may be present to enable the transformation of the microridge actin cytoskeleton conformation from a largely branched to a combination of branched and bundled actin, under certain physiological conditions. In fact, Bereiter-Hahn and co-workers have shown that microridges on the scale epidermis of adult guppy females exhibit a few actin bundles spaced along the length of the microridges (*Bereiter-Hahn et al.*, 1979).

During development, both the Arp2/3 complex as well as WASL localize to nascent microridges. In addition, the function of the Arp2/3 complex is essential for formation of actin punctae suggesting that they are formed of branched actin. Our data refutes the possibility that microridges are formed from microvilli (*Gorelik et al.*, 2003). In contrast to the Arp2/3 complex, which is associated with punctae even at 50% epiboly, keratin levels in the tissue are relatively low at early time points and rise concomitant with the growth of microridges. This increase in keratin levels might be linked to the specification of the periderm and could therefore be a trigger for microridge formation.

The ubiquitous presence of microridges during evolution suggests an important function for them. Microridges have been proposed to have various important functions, the most common one being the retention of mucus. Indeed, disruption of microridges by LatA, visibly affected the glycan layer. From both confocal and SEM data, it is clear that glycans are preferentially retained around the spot-like microridge remnants. Although, Arp2/3 complex inhibition had only a mild effect on the glycan layer, our TEM, SEM and WGA staining data as a whole suggests that glycans are enriched around microridges. WGA staining also follows the microridge pattern in the case of koi epidermal cells (*DePasquale*, 2018). It is thus likely that the distance between microridges is an important factor in the organization of the glycan layer, which is corroborated by the fact that the glycan layer on the head, which has closely packed microridges, exhibits a dense and uniform organization as compared to that on the flank, where microridges are spaced apart.

In summary, the actin microridge is a unique protrusion that has potential evolutionary origin from and structural homology to the leading edge of migrating cells, with an underlying actin and keratin containing sub-protrusive zone and functions in the organization of the overlying glycan layer. It will serve as an excellent model to further probe the role and regulation of actin networks in vivo.

## Materials and Methods

### Fish strains

For experiments, the zebrafish Tübingen (Tü) wild-type strain or GFP-negative individuals from the background of the *Tg(actb1:GFP-utrCH)* line (*Behrndt et al*., 2012) were used. For zebrafish maintenance and experimentation, the guidelines recommended by the Committee for the Purpose of Control and Supervision of Experiments on Animals (CPCSEA), Govt. of India, were followed.

### Plasmid injections

Plasmids, EGFP-EPLIN beta - a gift from Elizabeth Luna (Addgene plasmid # 40948) - and pcDNA3-myc-FLNa WT - a gift from John Blenis (Addgene plasmid # 8982)(*Woo et al*., 2004) - were purified using Invitrogen’s plasmid mini-prep kit and dissolved in autoclaved Milli-Q or nuclease free water. Plasmids were injected at a concentration of 40 ng/μl in autoclaved Milli-Q water at the 1 cell stage.

### Inhibitor treatments

CK666 (182515, Calbiochem), its inactive control CK689 (182517, Calbiochem) and Latrunculin A (L5163, Sigma) were dissolved in DMSO. For treatment with inhibitors, 10 dechorionated embryos/larvae were added in 1 ml E3 without methylene blue to a well of a 12 well plate and 1 ml of 2x inhibitor or vehicle in E3 without methylene blue was added to it and mixed gently with a pipette. CK666 and CK689 treatments (100 μM) were carried out for 1h whereas Latrunculin treatment (2 μM) was performed for 30 min.

### Immunohistology and imaging

For immunostainings, larvae were fixed in 4% PFA for 30mins and then overnight at 4°C. They were washed in PBS and permeabilized with PBS containing 0.8% Triton X-100 (PBSTx), blocked in 10% NGS in PBSTx for 3 h, incubated with primary antibody in 1% NGS in PBSTx for 3 h to overnight (O/N), washed with PBSTx 5 times for 30 min each, incubated with secondary antibody in 1% NGS in PBS for 3-4 h., followed by 5 washes in PBSTx each for 10 min. Samples were then fixed for 30 min or overnight in PFA, washed twice with PBS and upgraded in 30%, 50%, 70% glycerol in PBS.

For ArpC2, WASL and Cofilin stainings heat induced antigen retrieval was required. The samples were equilibrated in Tris-Cl (150mM, pH9.0, for ArpC2 and WASL) or Sodium Citrate buffer (10mM Sodium Citrate, 0.05% Tween-20, pH 6.0, for Cofilin) for 5 min. The samples were then incubated with fresh Tris-Cl or Sodium citrate buffer at 70° C for 20 min. The samples were allowed to reach room temperature, washed in PBSTx, and then processed for immunostaining from the blocking step as above.

For phalloidin staining, larvae were fixed in 4% PFA as before or for 3 h at RT. The larvae were then washed 5 times in PBS for 10 min each, incubated in 1:40 Phalloidin for 3-4 h followed by 5 washes in PBS for 10 min each and upgradation in glycerol in PBS. Phalloidin Rhodamine (R415) or Alexa Fluor 488-phalloidin (A12379) (Molecular probes) were used. When phalloidin and antibody staining were performed together, Phalloidin (1:400 in PBSTx) was used at the secondary antibody step.

For *α*-Tubulin staining, larvae were fixed in ice cold Dent’s fixative (80 % methanol, 20 % DMSO) on ice for 30 mins then at −20° C for 2 hours to O/N. Larvae were then downgraded in a methanol series, 100 %, 70 %, 50 %, 30 % methanol in PBS. They were then washed twice with PBS following which the normal method for immunostainings from the permeabilization step was followed.

Primary antibodies used in this study include Mouse anti-GFP (clone 12A6, DSHB; 1:100), Rabbit anti-GFP (TP401, Torrey Pines Biolabs; 1:200), Anti-Cofilin (SAB4300577, Sigma-Aldrich; 1:100), Anti-ArpC2 (HPA008352, Sigma Prestige; 1:100), Anti-Wasl (HPA005750, Sigma Prestige; 1:100), Anti-Myc (9E 10-s, DSHB, 9E 10 was deposited to the DSHB by Bishop, J.M; 1:100), Anti-pan Cytokeratin [AE1/AE3] (ab27988, Abcam; 1:100), Anti- *α*-Tubulin (T9026, Sigma; 1:100). The secondary antibodies and their dilutions were as follows: Anti-mouse Alexa 488-conjugated (A-11029, Invitrogen; 1:250), Anti-rabbit Alexa 488-conjugated (A11034, Invitrogen; 1:250), Anti-rabbit Alexa 546-conjugated (A-11035, Invitrogen; 1:250), Goat anti-Rabbit Cy3 (111-165-144, Jackson ImmunoResearch; 1:750), Goat anti-Rabbit Cy5 (111-175-144, Jackson ImmunoResearch; 1:750), Goat anti-Mouse Cy3 (115-165-146, Jackson ImmunoResearch; 1:750), Goat anti-Mouse Cy5 (115-175-146, Jackson ImmunoResearch; 1:750). DSHB-GFP-12A6 was deposited to the DSHB by DSHB (DSHB Hybridoma Product DSHB-GFP-12A6).

Imaging was carried out using the Zeiss LSM 510 Meta or Zeiss LSM 710 with EC Plan-Neofluar 40X/1.30 oil objective at 2x zoom or Plan-Apochromat 63x/1.40 oil objective at 1.5x zoom (Zeiss). 1024 x1024 image dimensions were used, with an averaging of 4.

For localization experiments, animals were either uninjected and untreated or in some sets, animals used were injected with the LifeAct-TagRFP plasmid (Ibidi; 60102) (*Riedl et al*., 2008) at the 1 cell stage or were treated with 1% DMSO and are indicated as (LifeAct injected) or (1% DMSO) respectively. Number of sets and total number of animals are indicated in brackets as (sets; animals). Images are representative for the following: At 48 hpf: ArpC2 (2; 10), ArpC2 (LifeAct-injected)(2; 10), WASL (1; 5), WASL (LifeAct-injected)(1; 4), Cofilin (1; 6), Cofilin (LifeAct-injected)(1; 8), Keratin (2; 8), *α*-Tubulin (1; 5), *α*-Tubulin (1% DMSO)(1; 5). At 50% Epiboly: ArpC2 (1; 5), WASL (1; 5). At 8-10 hpf: ArpC2 (LifeAct-injected)(2; 15), WASL (LifeAct-injected)(1; 5). For plasmid injections, images are representative for: Eplin-*β* (2; 10), Filamin-A (1; 3).

### Estimation of keratin intensity

Samples for keratin estimation were imaged on the Zeiss LSM 710, at 12-bit depth. Images were analyzed in FIJI (*Schindelin et al*., 2012). A circle with a diameter of 10 μm was made at a place in the cell at one slice below the ridge slice and its mean intensity was obtained. It was ensured that no circle had any point at saturation within it. These intensities were then plotted using Rstudio (*RStudio Team*, 2016).

### Electron Microscopy (EM) and EM-tomography

For EM, larvae at 48 hpf were fixed with two protocols. The first is based on Tilney and Tilney, (*Tilney and Tilney*, 1994), with slight modifications. Briefly, larvae were fixed for 30 minutes at 4°C in 1% OsO_4_ and 1% glutaraldehyde in 0.1 M Phosphate buffer (PB) at pH 6.2. They were washed in ice cold Milli-Q water (Millipore), 3 times for 5 minutes each then en bloc stained in 0.5% Uranyl acetate for 1 hour at room temperature (RT), dehydrated in 30%, 50%, 70%, 90%, 100%, 100%, 100% acetone series for 5 min each. The samples were then brought in acetone: Epon-Araldite (1:1) for 30 min followed by acetone: Epon-Araldite (1:2) for 30 min, overnight in Epon-Araldite and then placed in moulds and cured at 60 °C. For the other protocol, samples were fixed in 2.5% glutaraldehyde and 1% tannic acid in 0.1% PB at pH 7.2 for 45 min and then overnight in just 2.5% glutaraldehyde in the same buffer, washed in Milli-Q water 6 times, en block stained with 0.5% uranyl acetate for 1 hour at RT and then dehydrated and embedded in the same way as above.

For standard EM, 70-100 nm sections were collected on formvar coated slot or mesh grids or uncoated mesh grids and post stained with uranyl acetate and lead citrate and imaged on a Tecnai 12 microscope. For EM tomography, 300 nm sections were collected on formvar coated slot grids, further carbon coated, post stained with uranyl acetate, lead citrate and then with colloidal gold to add fiducials to both sides of the grid (10 nm gold nanoparticles, Sigma-Aldrich).

Tomograms were acquired and reconstructed as previously described (*Stepanek and Pigino*, 2016) with modifications. Electron tomograms were collected on a Tecnai F-30 (FEI) TEM at 300kV with a 2048×2048 Gatan CCD camera. Tilt series were collected in dual tilt axis geometry and maximum tilt range of 64°-65° and tilt steps of 1° in an automated manner using SerialEM software (*Mastronarde*, 2005). For the large tomogram (Fig. 1. A-I) a montage of 9 images was collected. The IMOD software package (*Kremer et al.*, 1996) was used to reconstruct and visualize the tomograms.

### EM tomogram segmentation

The EM images were de-noised using a low pass Gaussian smoothing followed by the Perona-Malik anisotropic diffusion method (*Perona and Malik*, 1990). These steps ensured image smoothing keeping region boundaries and small structures within the image unperturbed. Subsequently, to analyze the local behavior, 2D Hessian images were constructed as explained previously (*Bhavna et al.*, 2016). We exploited the Eigenvalues of the Hessian to extract actin structures within each image (*Frangi et al*., 1998). The higher of the Eigenvalues encompassed the salient features of the actin structures within the microridges. The eigenvalue images were binarised and similar pixels were connected in depth to obtain an n-dimensional image of the tomogram.

### Scanning Electron Microscopy on intact and detergent extracted samples

For detergent extraction, a previously published protocol (*Svitkina*, 2009) with a few modifications was used. Larvae were dechorionated and detergent extracted in 2 ml tubes with 500 μl PEM buffer (100 mM PIPES (free acid) pH 6.9, 1 mM MgCl2 and 1 mM EGTA) containing 2% PEG (M.wt. 35,000), 1% TritonX-100 and 2 μM phalloidin for 30 seconds. To the same tube, without removing any solution, 500 μl 0.4% glutaraldehyde in PEM containing 0.5% TritonX-100 was added and incubated for 2.5 min. Most of the solution was removed (at no point was the sample allowed to come in contact with the air-solution interface) and 1 ml of 2% TritonX-100 in PEM containing 2 μM phalloidin was added and the samples were incubated for 7 min followed by 3 washes with PEM containing 2 μM phalloidin for 1 min each. The samples were fixed in 2% glutaraldehyde in PEM (without phalloidin) for 20 min at RT and then O/N at 4 °C.

The samples were washed thrice with distilled water for 5 min each, treated with a 0.2% tannic acid solution in distilled water for 20 min, washed again 5 times with distilled water for 5 min each and then treated with 0.4% uranyl acetate in distilled water for 1 h at RT. Samples were then washed 3 times with distilled water and dehydrated in an acetone series (30%, 50%, 60%, 70%, 80%, 90%, 100%, 100%, 100% for 5 min each) and 1:1 Hexamethyldisilazane (440191, Sigma): Acetone for 5 min. Subsequently, neat Hexamethyldisilazane treatment was carried out for 5 min and again for 10 min followed by removal of the excess Hexamethyldisilazane. The samples were picked with filter paper and left to air dry for 2-4 h. Post drying samples were placed on a stub with carbon tape, sputter coated with gold for 1 min and imaged with a Zeiss Gemini SEM. For detergent extracted samples, SEM was performed on 3 animals in total from 2 sets.

For SEM on non-detergent extracted larvae fixation was done in 2.5% gluteraldehyde for 30 min at RT and then overnight at 4 °C. Then either the protocol described above for detergent extraction samples from the washes and tannic acid step or an OTOTO protocol was followed with changes (*Fischer et al*., 2012). For OTOTO, the samples were first stained with OsO_4_ (O) for 15 min, washed 3 times, stained with saturated aqueous Thiocarbohydrazide (T) (T2137, Sigma) for 15 min and washed 3 times. This was repeated till the O-T-O-T-O steps were completed and then dehydrated with acetone as above for the detergent extracted samples, without sputter coating.

### Preservation and imaging of the glycan layer

For imaging by light microscopy, larvae at 48 hpf were fixed with 75 mM Lysine (124828, SRL), 0.1M Cacodylate buffer (11655, EMS), 2% PFA (103999, Merck), 2.5% gluteraldehyde (11614, EMS). The PFA and gluteraldehyde were added together just prior to use. The larvae were fixed for 22-24 h at RT. The samples were then washed 4-5 times with PBS. They were then incubated with WGA-594 (W11262, Invitrogen; 1:200) along with Phalloidin-Alexa488 in PBS for 3-4 h at RT, followed by 5 washes with PBS (5-10 min each). Finally, the samples were serially upgraded into 70% glycerol and mounted for confocal microscopy.

For imaging by SEM, a protocol similar to (*Fischer et al*., 2012) was used. Larvae at 48 hpf were fixed with a cocktail containing 75 mM Lysine (124828, SRL), 0.1M Cacodylate buffer (11655, EMS), 2% PFA (103999, Merck), 2.5% gluteraldehyde (11614, EMS), 0.075% Alcian Blue (48261, SRL) for 22-24 h at RT. Then the protocol for OTOTO staining was followed including the dehydration and imaging steps.

For TEM, the above SEM protocol was used for sample preparation, including the OTOTO steps, then the samples were dehydrated, embedded and processed for EM analysis as above and imaged in a Zeiss Libra 120 microscope at 120 kV. Images are representative for the following: SEM (2 sets; 6 animals in total), WGA (2 sets; 12 animals in total)

### Estimation of microridge parameters

For estimation of microridge height, microridges that appeared to be straight were measured from their base to the tip manually in FIJI using TEM images. For estimation of microridge width, measurements were made roughly perpendicular to the microridge length manually in FIJI using SEM images, some microridges were estimated at multiple locations along their length.

### Statistical analysis and image processing

Processing of fluorescence microscopy images was carried out using FIJI. Panels were made in ScientiFig (*Aigouy and Mirouse*, 2013) and exported to Adobe Illustrator or directly made in Inkscape, statistical analysis and plot generation (using the ggplot2 package) was carried out using R in Rstudio (*RStudio Team*, 2016; *Wickham*, 2016; *R Core Team*, 2018). Description of the boxplots — the line in the middle is the median, lower and upper box limits show the 1st quartile (Q1) and 3rd quartile (Q3) positions respectively. The whiskers extend 1.5 times the interquartile range or to the furthest point from the box limit, whichever is closer to the box limit and each point indicates a data point used in generating the box-plot (*Spitzer et al*., 2014). Images for the model were created in Blender (Blender Foundation).

## Acknowledgments

We acknowledge Pushan Ayyub, Bhagyashri Chalke, Rudheer Bapat, Seema Shirolikar, Lalit Borde and Tobias Fürstenhaupt for help in SEM/TEM and TIFR and Wellcome-trust-DBT-India-Alliance (MS) for funding. The authors have no additional competing financial interests. We also thank the Henriques Lab and eLIFE for their Latex templates that we modified and used on Overleaf to generate this manuscript.

## Description of Videos

**Video 1.** An electron tomogram through the long axis of the microridge. Each frame represents a 0.7 nm thick section in the Z axis. The scale bar is 200 nm. The sample was prepared using glutaraldehyde and OsO_4_. This is the same tomogram shown in Figure 1 C-H.

**Video 2.** An electron tomogram of a transverse section of a microridge at the cell junction. Each frame represents a 0.55 nm thick section in the Z axis. The scale bar is 200 nm. The sample was prepared using glutaraldehyde and OsO_4_. This is the same tomogram shown in Figure 1 I-K.

**Video 3.** An electron tomogram at an angle through the transverse section of the microridge. Each frame represents a 0.55 nm thick section in the Z axis. The sample was prepared using glutaraldehyde and tannic acid but without OsO_4_. The tannic acid makes actin filaments appear thicker than in the previous cases. The scale bar is 200 nm.

**Video 4.** Segmentation of an electron tomogram of a microridge reveals a network of actin. A segmented region shown in Figure 2 B,F of the tomogram in 3D. The Z-depth is indicated by the colourbar.

## Supplementary Figures

**Fig. S 1.**
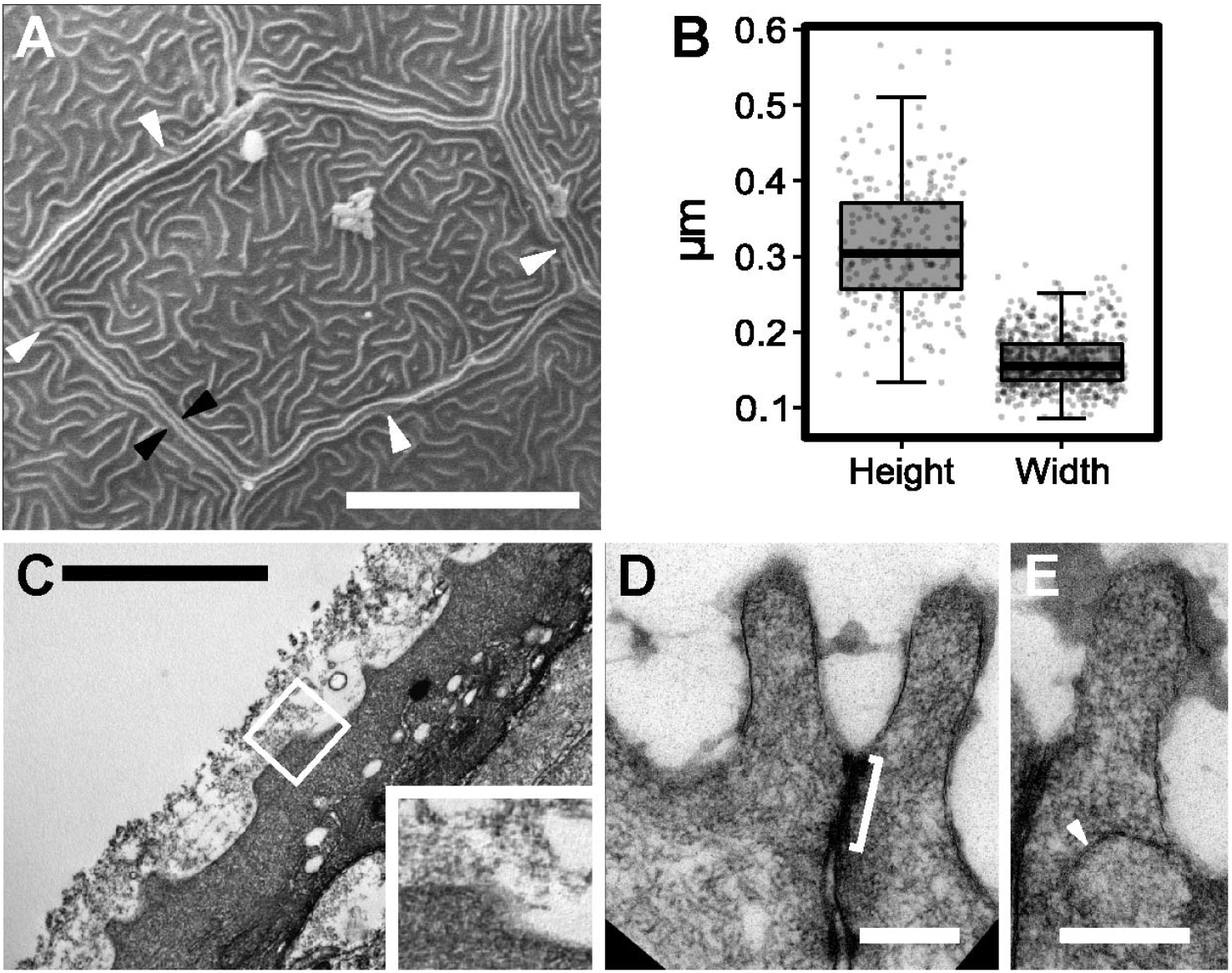
Ultrastructural analysis of the peridermal apical domain. Scanning electron micrographs of the head periderm (A). SEM analysis shows that long microridges (black arrowheads in A) flank the tight junctions on either side of the cell margin. These junctional microridges may be discontinuous (white arrows in A). Quantification of microridge height and width (B) as measured by TEM and SEM, respectively. TEM (C) analysis in presence of Alcian Blue and Lysine revealed a thick layer of mucus present above microridges. Inset in C, which is high zoom of the boxed region, shows mucus above and around the microridge. Normal TEM of microridges (D,E). A tight junction as shown by the bracket in D connects the two cells. Note the presence of vesicles at the location shown by the white arrowhead in E. Images obtained from 48 hpf larvae. Scale bar in A represents 10 μm, in C is 2 μm and in D,E are equal to 0.2 μm.

**Fig. S 2.**
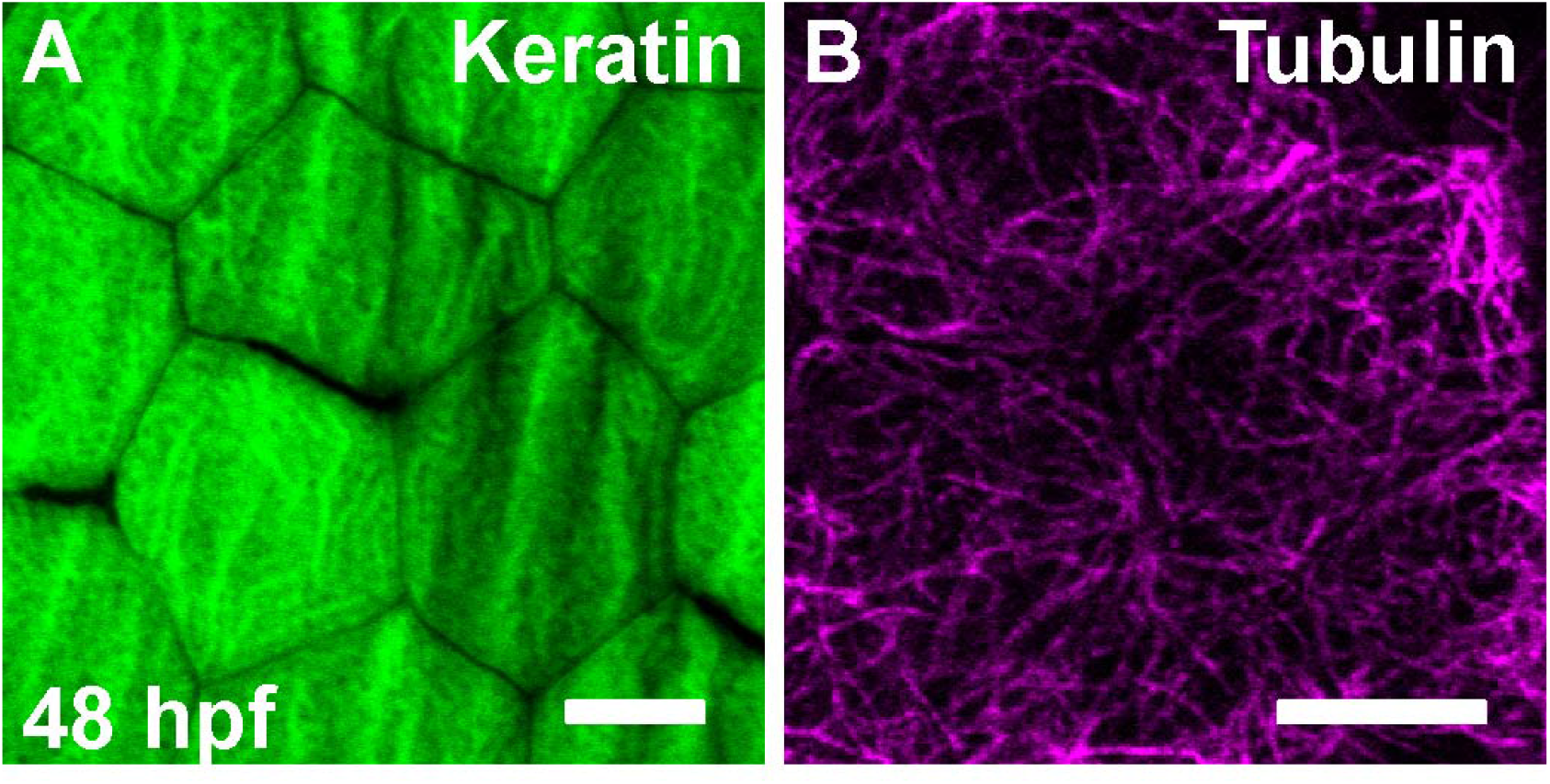
The localization of keratin and microtubules at the peridermal apical and sub-apical domains. Confocal microscopy analysis of peridermal cells showing localization of apical keratin (A), and sub-apical microtubules (B) at 48 hpf. Note that keratin follows the microridge pattern in parts but microtubules do not. Images obtained from 48 hpf larvae. Scale bars in A,B represent 10 μm.

**Fig. S 3.**
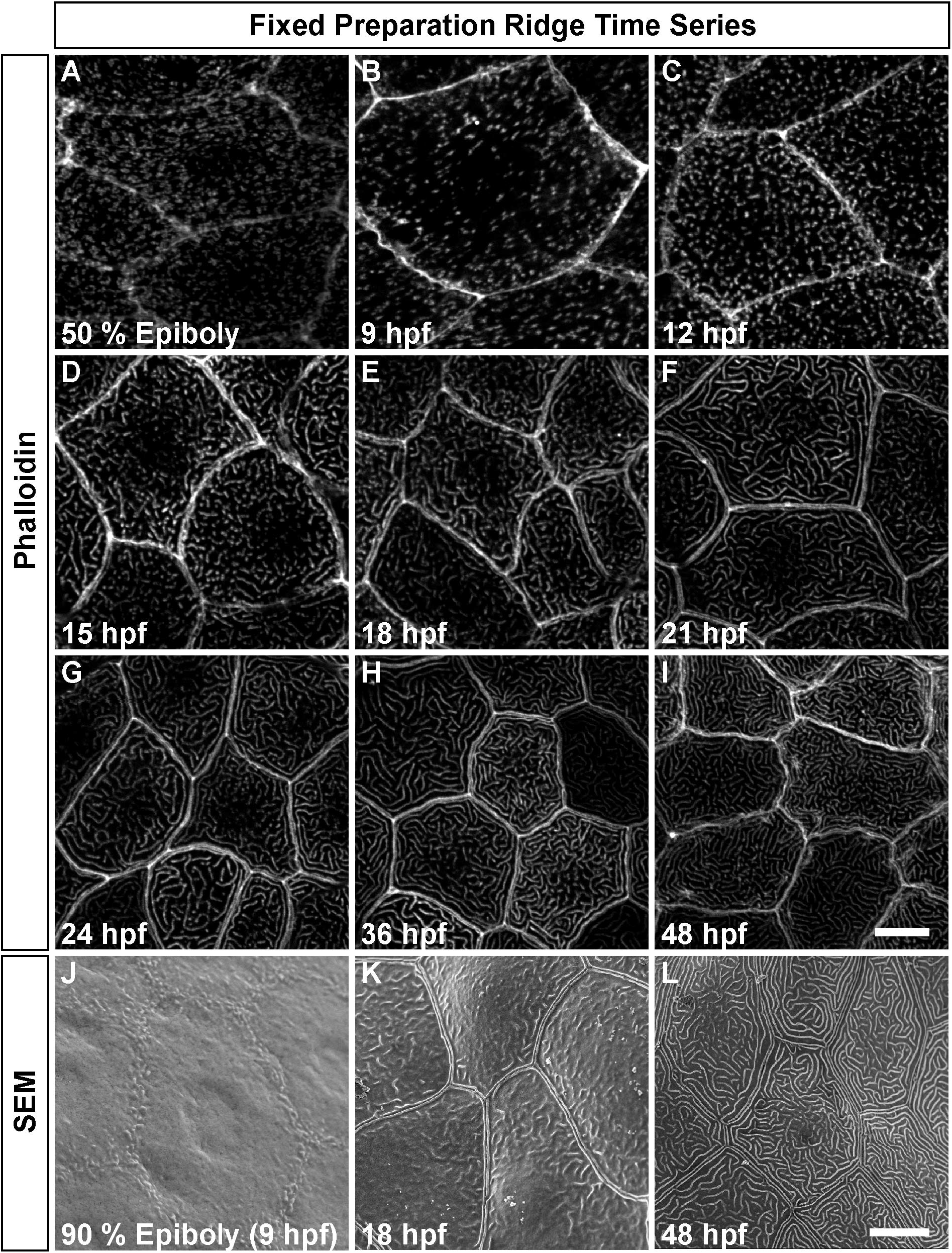
Formation of microridges during early embryogenesis. Confocal microscopy analysis of developing microridges by phalloidin staining (A-I) and SEM (J-L) at the given stages. Note that at 9 hpf the actin punctae do not form projections (B,J). The microridges begin to form at the cell periphery at 18 hpf and their length increases over time. The scale bars in I for A-I and in L for J-L are 10 μm. The images are representative for: Phalloidin stainings: a total of 10 animals from 2 sets; SEM: a total of 6 animals from 2 sets.

**Fig. S 4.**
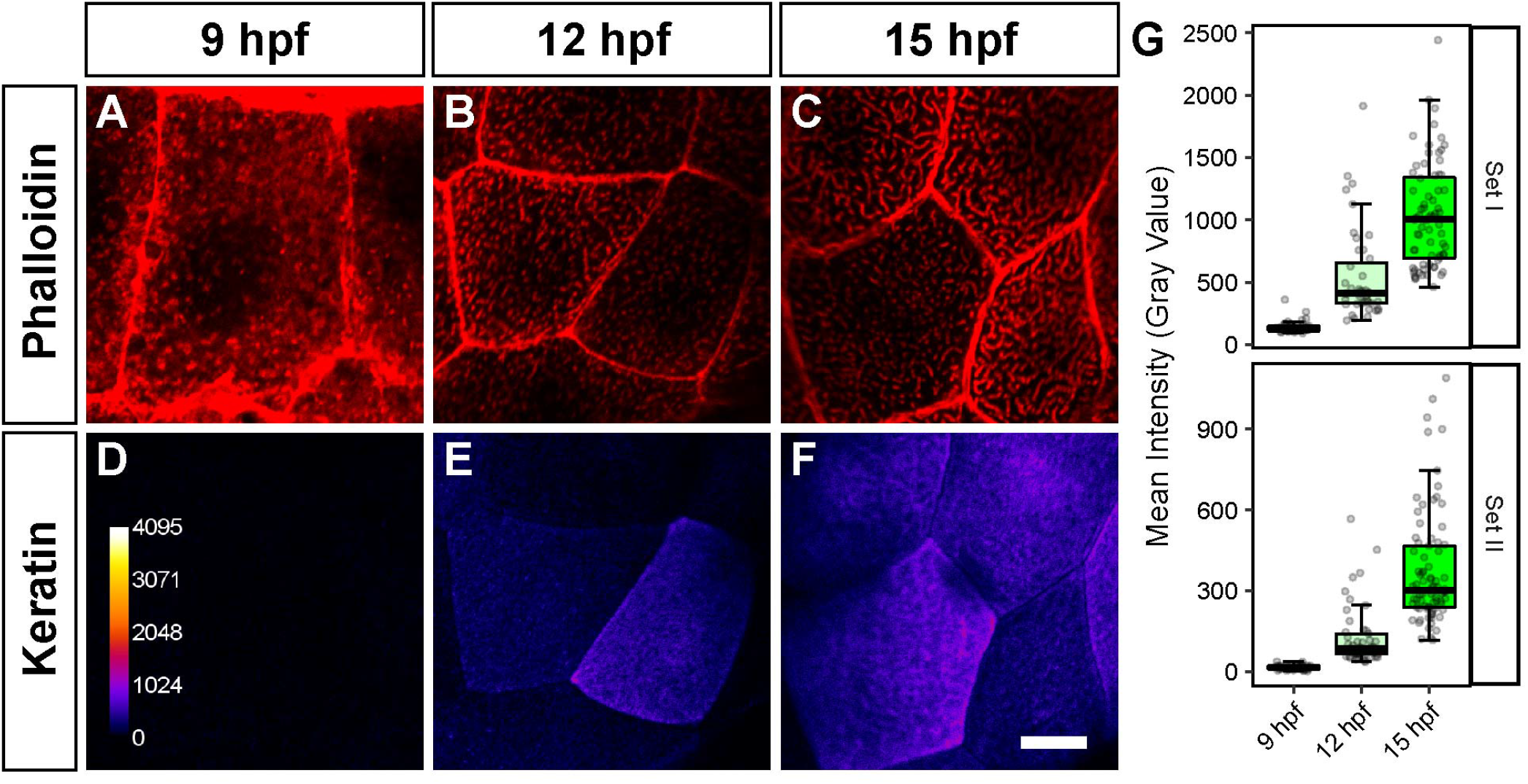
Keratin intensity during early development. (A-C) Phalloidin staining at 9 (A), 12 (B), and 15 (C) hpf in red and corresponding keratin stainings in fire pseudocolour (D-F). Boxplots (G) showing keratin intensities for two separate experimental sets. There were 4-6 animals per set. The calibration bar for the keratin staining intensity in D is for D-F. Scale bar in F for A-F represents 10 μm.

## References

Abe K, Takeichi M. EPLIN mediates linkage of the cad-herin catenin complex to F-actin and stabilizes the circumferential actin belt. Proc Natl Acad Sci U S A. 2008; 105(1):13–9. https://www.ncbi.nlm.nih.gov/pubmed/18093941, doi: 10.1073/pnas.0710504105.

Aigouy B, Mirouse V. ScientiFig: a tool to build publication-ready scientific figures. Nat Methods. 2013; 10(11):1048. https://www.ncbi.nlm.nih.gov/pubmed/24173380, doi: 10.1038/nmeth.2692.

Andrews PM. Microplicae: characteristic ridge-like folds of the plasmalemma. J Cell Biol. 1976; 68(3):420–9. https://www.ncbi.nlm.nih.gov/pubmed/828906.

Apodaca G, Gallo LI. Epithelial Polarity. Colloquium Series on Building Blocks of the Cell: Cell Structure and Function. 2013 mar; 1(2):1–115. http://www.morganclaypool.com/doi/abs/10.4199/C00077ED1V01Y201303BBC002, doi: 10.4199/C00077ED1V01Y201303BBC002.

Baskin JM, Dehnert KW, Laughlin ST, Amacher SL, Bertozzi CR. Visualizing enveloping layer glycans during zebrafish early embryogenesis. Proc Natl Acad Sci U S A. 2010; 107(23):10360–5. https://www.ncbi.nlm.nih.gov/pubmed/20489181, doi: 10.1073/pnas.0912081107.

Behrndt M, Salbreux G, Campinho P, Hauschild R, Oswald F, Roensch J, Grill SW, Heisenberg CP. Forces Driving Epithelial Spreading in Zebrafish Gastrulation. Science. 2012; 338(6104):257–260. http://science.sciencemag.org/content/338/6104/257, doi: 10.1126/science.1224143.

Bereiter-Hahn J, Osborn M, Weber K, Voth M. Filament organization and formation of microridges at the surface of fish epidermis. J Ultrastruct Res. 1979; 69(3):316–30. https://www.ncbi.nlm.nih.gov/pubmed/159961.

Bhavna R, Uriu K, Valentin G, Tinevez JY, Oates AC. Correction: Object Segmentation and Ground Truth in 3D Embryonic Imaging. PLoS One. 2016; 11(8):e0161550. https://www.ncbi.nlm.nih.gov/pubmed/27529424, doi: 10.1371/journal.pone.0161550.

Breipohl W, Herberhold C, Kerschek R. Microridge cells in the larynx of the male white rat. Investigations by reflection scanning electron microscopy. Arch Otorhinolaryngol. 1977; 215(1):1–9. https://www.ncbi.nlm.nih.gov/pubmed/577135.

Burnette DT, Manley S, Sengupta P, Sougrat R, Davidson MW, Kachar B, Lippincott-Schwartz J. A role for actin arcs in the leading-edge advance of migrating cells. Nat Cell Biol. 2011; 13(4):371–81. http://www.ncbi.nlm.nih.gov/pubmed/21423177, doi: 10.1038/ncb2205.

Collin SP, Collin HB. A comparative SEM study of the vertebrate corneal epithelium. Cornea. 2000; 19(2):218–30. https://www.ncbi.nlm.nih.gov/pubmed/10746456.

Cone RA. Barrier properties of mucus. Adv Drug Deliv Rev. 2009; 61(2):75–85. https://www.ncbi.nlm.nih.gov/pubmed/19135107, doi: 10.1016/j.addr.2008.09.008.

Crawley SW, Mooseker MS, Tyska MJ. Shaping the intestinal brush border. J Cell Biol. 2014; 207(4):441–51. https://www.ncbi.nlm.nih.gov/pubmed/25422372, doi: 10.1083/jcb.201407015.

DePasquale JA. Microridges in Cyprinus carpio scale epidermis. Acta Zoologica. 2018; 99(2):158–168. https://onlinelibrary.wiley.com/doi/abs/10.1111/azo.12201, doi: doi:10.1111/azo.12201.

Eroschenko VP, Osman F. Scanning electron microscopic changes in vaginal epithelium of suckling neonatal mice in response to estradiol or insecticide chlordecone (Kepone) passage in milk. Toxicology. 1986; 38(2):175–85. https://www.ncbi.nlm.nih.gov/pubmed/2418535.

Fassel TA, Mozdziak PE, Sanger JR, Edmiston CE. Paraformaldehyde effect on ruthenium red and lysine preservation and staining of the staphylococcal glycocalyx. Microsc Res Tech. 1997; 36(5):422–7. https://www.ncbi.nlm.nih.gov/pubmed/9140944, doi: 10.1002/(SICI)1097-0029(19970301)36:5<422::AID-JEMT12>3.0.CO;2-U.

Fassel TA, Schaller MJ, Remsen CC. Comparison of alcian blue and ruthenium red effects on preservation of outer envelope ultrastructure in methanotrophic bacteria. Microsc Res Tech. 1992; 20(1):87–94. https://www.ncbi.nlm.nih.gov/pubmed/1377060, doi: 10.1002/jemt.1070200109.

Fenger C, Knoth M. The anal transitional zone: a scanning and transmission electron microscopic investigation of the surface epithelium. Ultrastruct Pathol. 1981; 2(2):163–73. https://www.ncbi.nlm.nih.gov/pubmed/7268926.

Fischer ER, Hansen BT, Nair V, Hoyt FH, Dorward DW. Scanning electron microscopy. Curr Protoc Microbiol. 2012; Chapter 2:Unit 2B 2. https://www.ncbi.nlm.nih.gov/pubmed/22549162, doi: 10.1002/9780471729259.mc02b02s25.

Fishelson L. A comparative study of ridge-mazes on surface epithelial cell-membranes of fish scales (Pisces, Teleostei). Zoomorphology. 1984; 104(4):231–238. https://doi.org/10.1007/BF00312036, doi: 10.1007/bf00312036.

Frangi AF, Niessen WJ, Vincken KL, Viergever MA. Muliscale Vessel Enhancement Filtering. In: Proceedings of the First International Conference on Medical Image Computing and Computer-Assisted Intervention MICCAI ‘98, London, UK, UK: Springer-Verlag; 1998. p. 130–137. http://dl.acm.org/citation.cfm?id=646921.709471.

Gorelik J, Shevchuk AI, Frolenkov GI, Diakonov IA, Lab MJ, Kros CJ, Richardson GP, Vodyanoy I, Edwards CRW, Klenerman D, Korchev YE. Dynamic assembly of surface structures in living cells. Proceedings of the National Academy of Sciences. 2003; 100(10):5819–5822. http://www.pnas.org/content/100/10/5819, doi: 10.1073/pnas.1030502100.

Hafez ES, Fadel HE, Noonan SM, Oshima M, Okamura H, Watson JH, Zaneveld LJ, Steger RW. Scanning electron microscopy of human female reproductive tract and amniotic fluid cells. Int J Fertil. 1977; 22(4):193–205. https://www.ncbi.nlm.nih.gov/pubmed/24598.

Hawkes JW. The structure of fish skin. I. General organization. Cell Tissue Res. 1974; 149(2):147–58. https://www.ncbi.nlm.nih.gov/pubmed/4424315.

Johnston BT, Nunn S, Sloan JM, Collins JS, McFarland RJ, Parkin S, Carr KE, Collins BJ. The application of microridge analysis in the diagnosis of gastro-oesophageal reflux disease. Scand J Gastroenterol. 1996; 31(2):97–102. https://www.ncbi.nlm.nih.gov/pubmed/8658046.

Kimmel CB, Warga RM, Schilling TF. Origin and organization of the zebrafish fate map. Development. 1990; 108(4):581–94. https://www.ncbi.nlm.nih.gov/pubmed/2387237.

Kremer JR, Mastronarde DN, McIntosh JR. Computer visualization of three-dimensional image data using IMOD. J Struct Biol. 1996; 116(1):71–6. https://www.ncbi.nlm.nih.gov/pubmed/8742726, doi: 10.1006/jsbi.1996.0013.

Lam PY, Mangos S, Green JM, Reiser J, Huttenlocher A. In vivo imaging and characterization of actin microridges. PLoS One. 2015; 10(1):e0115639. http://www.ncbi.nlm.nih.gov/pubmed/25629723, doi: 10.1371/journal.pone.0115639.

Le Bivic A. Evolution and cell physiology. 4. Why invent yet another protein complex to build junctions in epithelial cells? Am J Physiol Cell Physiol. 2013; 305(12):C1193–201. https://www.ncbi.nlm.nih.gov/pubmed/24025867, doi: 10.1152/ajpcell.00272.2013.

Makabe S, Motta PM, Naguro T, Vizza E, Perrone G, Zichella L. Microanatomy of the female reproductive organs in postmenopause by scanning electron microscopy. Climacteric. 1998; 1(1):63–71. https://www.ncbi.nlm.nih.gov/pubmed/11907929.

Mastronarde DN. Automated electron microscope tomography using robust prediction of specimen movements. Journal of Structural Biology. 2005; 152(1):36–51. http://www.sciencedirect.com/science/article/pii/S1047847705001528, doi: https://doi.org/10.1016/jjsb.2005.07.007.

Maul RS, Song Y, Amann KJ, Gerbin SC, Pollard TD, Chang DD. EPLIN regulates actin dynamics by crosslinking and stabilizing filaments. J Cell Biol. 2003; 160(3):399–407. https://www.ncbi.nlm.nih.gov/pubmed/12566430, doi: 10.1083/jcb.200212057.

Perona P, Malik J. Scale-space and edge detection using anisotropic diffusion. IEEE Transactions on Pattern Analysis and Machine Intelligence. 1990; 12(7):629–639. doi: 10.1109/34.56205.

R Core Team. R: A Language and Environment for Statistical Computing. R Foundation for Statistical Computing, Vienna, Austria; 2018, https://www.R-project.org/.

Raman R, Damle I, Rote R, Banerjee S, Dingare C, Sonawane M. aPKC regulates apical localization of Lgl to restrict elongation of microridges in developing zebrafish epidermis. Nat Commun. 2016; 7:11643. https://www.ncbi.nlm.nih.gov/pubmed/27249668, doi: 10.1038/ncomms11643.

Revenu C, Athman R, Robine S, Louvard D. The co-workers of actin filaments: from cell structures to signals. Nat Rev Mol Cell Biol. 2004; 5(8):635–46. http://www.ncbi.nlm.nih.gov/pubmed/15366707, doi: 10.1038/nrm1437.

Riedl J, Crevenna AH, Kessenbrock K, Yu JH, Neukirchen D, Bista M, Bradke F, Jenne D, Holak TA, Werb Z, Sixt M, Wedlich-Soldner R. Lifeact: a versatile marker to visualize F-actin. Nature Methods. 2008; 5:605. http://dx.doi.org/10.1038/nmeth.1220, doi: 10.1038/nmeth.1220.

Rodriguez-Boulan E, Nelson WJ. Morphogenesis of the polarized epithelial cell phenotype. Science. 1989; 245(4919):718–25. https://www.ncbi.nlm.nih.gov/pubmed/2672330.

RStudio Team. RStudio: Integrated Development Environment for R. RStudio, Inc., Boston, MA; 2016, http://www.rstudio.com/.

Saito H, Itoh I. Ultrastructural study of rabbit buccal epithelial cells and intercellular junction by scanning and transmission electron microscopy. J Electron Microsc (Tokyo). 1993; 42(6):389–93. https://www.ncbi.nlm.nih.gov/pubmed/8176332.

Sauvanet C, Wayt J, Pelaseyed T, Bretscher A. Structure, regulation, and functional diversity of microvilli on the apical domain of epithelial cells. Annu Rev Cell Dev Biol. 2015; 31:593–621. https://www.ncbi.nlm.nih.gov/pubmed/26566117, doi: 10.1146/annurev-cellbio-100814-125234.

Schindelin J, Arganda-Carreras I, Frise E, Kaynig V, Longair M, Pietzsch T, Preibisch S, Rueden C, Saalfeld S, Schmid B, Tinevez JY, White DJ, Hartenstein V, Eliceiri K, Tomancak P, Cardona A. Fiji: an open-source platform for biological-image analysis. Nat Methods. 2012; 9(7):676–82. https://www.ncbi.nlm.nih.gov/pubmed/22743772, doi: 10.1038/nmeth.2019.

Schliwa M. Cytoarchitecture of surface layer cells of the teleost epidermis. Journal of Ultrastructure Research. 1975;.

Sharma A, Anderson KI, Muller DJ. Actin microridges characterized by laser scanning confocal and atomic force microscopy. FEBS Lett. 2005; 579(9):2001–8. http://www.ncbi.nlm.nih.gov/pubmed/15792810, doi: 10.1016/j.febslet.2005.02.049.

Small JV, Stradal T, Vignal E, Rottner K. The lamellipodium: where motility begins. Trends Cell Biol. 2002; 12(3):112–20. https://www.ncbi.nlm.nih.gov/pubmed/11859023.

Sperry DG, Wassersug RJ. A proposed function for microridges on epithelial cells. Anat Rec. 1976; 185(2):253–7. https://www.ncbi.nlm.nih.gov/pubmed/1275311, doi: 10.1002/ar.1091850212.

Spitzer M, Wildenhain J, Rappsilber J, Tyers M. BoxPlotR: a web tool for generation of box plots. Nat Methods. 2014; 11(2):121–2. https://www.ncbi.nlm.nih.gov/pubmed/24481215, doi: 10.1038/nmeth.2811.

Stepanek L, Pigino G. Microtubule doublets are double-track railways for intraflagellar transport trains. Science. 2016; 352(6286):721–724. https://science.sciencemag.org/content/352/6286/721, doi: 10.1126/science.aaf4594.

Svitkina T. Imaging cytoskeleton components by electron microscopy. Methods Mol Biol. 2009; 586:187–206. https://www.ncbi.nlm.nih.gov/pubmed/19768431, doi: 10.1007/978-1-60761-376-3_10.

Svitkina TM, Borisy GG. Arp2/3 complex and actin depolymerizing factor/cofilin in dendritic organization and tread-milling of actin filament array in lamellipodia. J Cell Biol. 1999; 145(5):1009–26. https://www.ncbi.nlm.nih.gov/pubmed/10352018.

Tilney LG, Derosier DJ, Mulroy MJ. The organization of actin filaments in the stereocilia of cochlear hair cells. J Cell Biol. 1980; 86(1):244–59. https://www.ncbi.nlm.nih.gov/pubmed/6893452.

Tilney LG, Tilney MS. Methods to visualize actin polymerization associated with bacterial invasion. In: Bacterial Pathogenesis Part B: Interaction of Pathogenic Bacteria with Host Cells, vol. 236 of Methods in Enzymology Academic Press; 1994.p. 476–481. http://www.sciencedirect.com/science/article/pii/0076687994360367, doi: https://doi.org/10.1016/0076-6879(94)36036-7.

Uehara K, Miyoshi M, Miyoshi S. Cytoskeleton in microridges of the oral mucosal epithelium in the carp, Cyprinus carpio. Anat Rec. 1991; 230(2):164–8. https://www.ncbi.nlm.nih.gov/pubmed/1714256, doi: 10.1002/ar.1092300203.

Urdy S, Goudemand N, Pantalacci S. Looking Beyond the Genes: The Interplay Between Signaling Pathways and Mechanics in the Shaping and Diversification of Epithelial Tissues. Curr Top Dev Biol. 2016; 119:227–90. https://www.ncbi.nlm.nih.gov/pubmed/27282028, doi: 10.1016/bs.ctdb.2016.03.005.

Vadlamudi RK, Li F, Adam L, Nguyen D, Ohta Y, Stossel TP, Kumar R. Filamin is essential in actin cytoskeletal assembly mediated by p21-activated kinase 1. Nat Cell Biol. 2002; 4(9):681–90. https://www.ncbi.nlm.nih.gov/pubmed/12198493, doi: 10.1038/ncb838.

Wickham H. ggplot2: Elegant Graphics for Data Analysis. Springer-Verlag New York; 2016. http://ggplot2.org.

Woo MS, Ohta Y, Rabinovitz I, Stossel TP, Blenis J. Ribosomal S6 Kinase (RSK) Regulates Phosphorylation of Filamin A on an Important Regulatory Site. Molecular and Cellular Biology. 2004; 24(7):3025–3035. http://mcb.asm.org/content/24/7/3025.abstract, doi: 10.1128/mcb.24.7.3025-3035.2004.

Worawongvasu R. A scanning electron microscopic study of the dysplastic epithelia adjacent to oral squamous cell carcinoma. Ultrastruct Pathol. 2007; 31(4):273–81. https://www.ncbi.nlm.nih.gov/pubmed/17786828, doi: 10.1080/01913120701515637.

